# Is obesity a genetic disease? Human obese transcriptome analysis in monozygotic twins

**DOI:** 10.1101/159186

**Authors:** Francesc Font-Clos, Stefano Zapperi, Caterina A.M. La Porta

## Abstract

Obesity is a pandemic disease with a critical increase in childhood. An important unanswered question is to understand if this disease is due to genetic causes or to the life-style of the subjects. To address this question, we have analyzed if monozygotic twins show the same robust transcriptomic signature (5σ, as for the Higgs Boson) that we have recently revealed in obese subjects. Our results show that our signature correlates with BMI in paired transcriptomes of monozygotic twins, suggesting that the signature does not reflect underlying genetic causes.

## Introduction

Obesity is a pandemic disease with an impressive increase in children.^1^ The contribution to obesity of a genetic background is still debated. Well established cases of Mendelian forms of obesity approximately account for only 5*%* of the severely obese cases.^2^ In the case of common obesity, recent genome wide association studies (GWAS) have investigated possible relations between single nucleotide polymorphism (SNP) and Body Mass Index (BMI).^3^ Despite the sheer amount of data and the effort devoted to this task, none of the resulting genetic loci have real predictive power. In particular, genetic contributions do not account for most BMI variations between subjects which are likely due to lifestyle and environmental factors.^3^ A recent paper reported the gene expression profile in subcutaneous adipose tissue of BMI-discordant monozygotic twin pairs without finding any molecular or clinical changes associated with subtypes of obesity.^4^

Recently, we revealed a robust transcriptomic signature of obesity by analyzing a collection of available datasets from adipocytes of obese subjects.^5^ We were able to obtain a strong statistical significance (5σ as for the discovery of the Higgs boson) by eliminating batch effects due to the mixing of different datasets. The signature comprises 38 genes involved in the interaction between cells and the extracellular matrix, inflammation and central nervous system.^5^

In this brief note, we investigate if a Mendelian contribution to obesity is relevant for our signature. To this end, we check if our signature is correlated with BMI in paired transcriptomes of monozygotic twins.

## Results

### The transcriptomic signature of obesity

Here we give a brief overview of the derivation of the transcriptomic signature of obesity, inviting the interested reader to see^5^ for details. Our analysis revolved around *SVDmerge* (https://github.com/ComplexityBiosystems/SVDmerge), an algorithm to remove batch effects, and *Pathway Deregulation Scores,* a pathway-based dimensionality reduction technique.^6^ Combining these two methodologies allowed us to (i) merge several publicly available datasets, increasing the number of samples in the analysis, and (ii) transition from a gene-based to a pathway-based perspective, decreasing the number of variables from ∼ 20000 genes to ∼ 1000 pathways. In this way we substantially improved the samples-to-variables ratio and were able to identify pathways related to adhesion molecules, inflammation, salivary secretion and digestive problems. We also proposed a simple obesity score, computed as a linear combination of the expression of the 38 genes, and showed that it correlates well with BMI in several independent validation datasets. We verified that such correlations are gender-independent and tissue-specific. Finally, we pointed out that some of the deregulation patterns found in obesity are also seen in breast tumor samples.

It is interesting to compare our transcriptomic signature with existing results on obesity based on GWAS.^3^ These studied have revealed a set of genetic loci that are associated with BMI variations. We have compared the list of genes in our signature with the list of genes reported in Ref.^3^ as significantly associated with BMI. The two lists have no intersection. Similarly, the list of significant pathways revealed in Ref.^3^ has no intersection with the list reported in.^5^ Therefore, our approach allows to identify genes that normally are not highlighted because we are able to analyze more datasets due to the removal of batch effects by the method of single value decomposition.^5^ The power of a Big Data analysis is actually to uncover things that are not easy to see, in this case genes and pathways at the roots of the problem.

## Analysis of transcriptomes from BMI-discordant monozygotic twins

We applied the same strategy described in the previous section to study transcriptomic data from monozygotic (MZ) twins with discordant BMI, see Methods for details. Figure 1 shows that the obesity score correlates with BMI (*R* = 0.68, *p* = 1.40 × 10^−4^) considering all the 26 samples of the batch. This data set is particularly interesting because it consists of 26 samples from 13 MZ twin pairs whose BMI is highly discordant. Because MZ twins are genetically identical, BMI variations between a subject and its co-twin should be due exclusively to environmental factors and lifestyle. Figure 1b shows indeed that the variations in BMI correlate with variations in score (*R* = 0.58,*p* = 3.80 × 10^−2^) when considering only pairing between co-twins. We then perform a randomization test and compare changes in BMI and score in sets of 13 randomly chosen pairs of unrelated subjects. As shown in Figure 1c, there is no significant difference (*p* = 0.35) between the correlation in co-twins and the one in unrelated twins. Hence, the signature in co-twins reflects merely the BMI, rather than the genetic background that should be identical in co-twins and different in randomly paired subjects. This suggests that our transcriptomic signature is associated with obesity rather than with any underlying genetic differences in the subjects.

**Figure 1.**
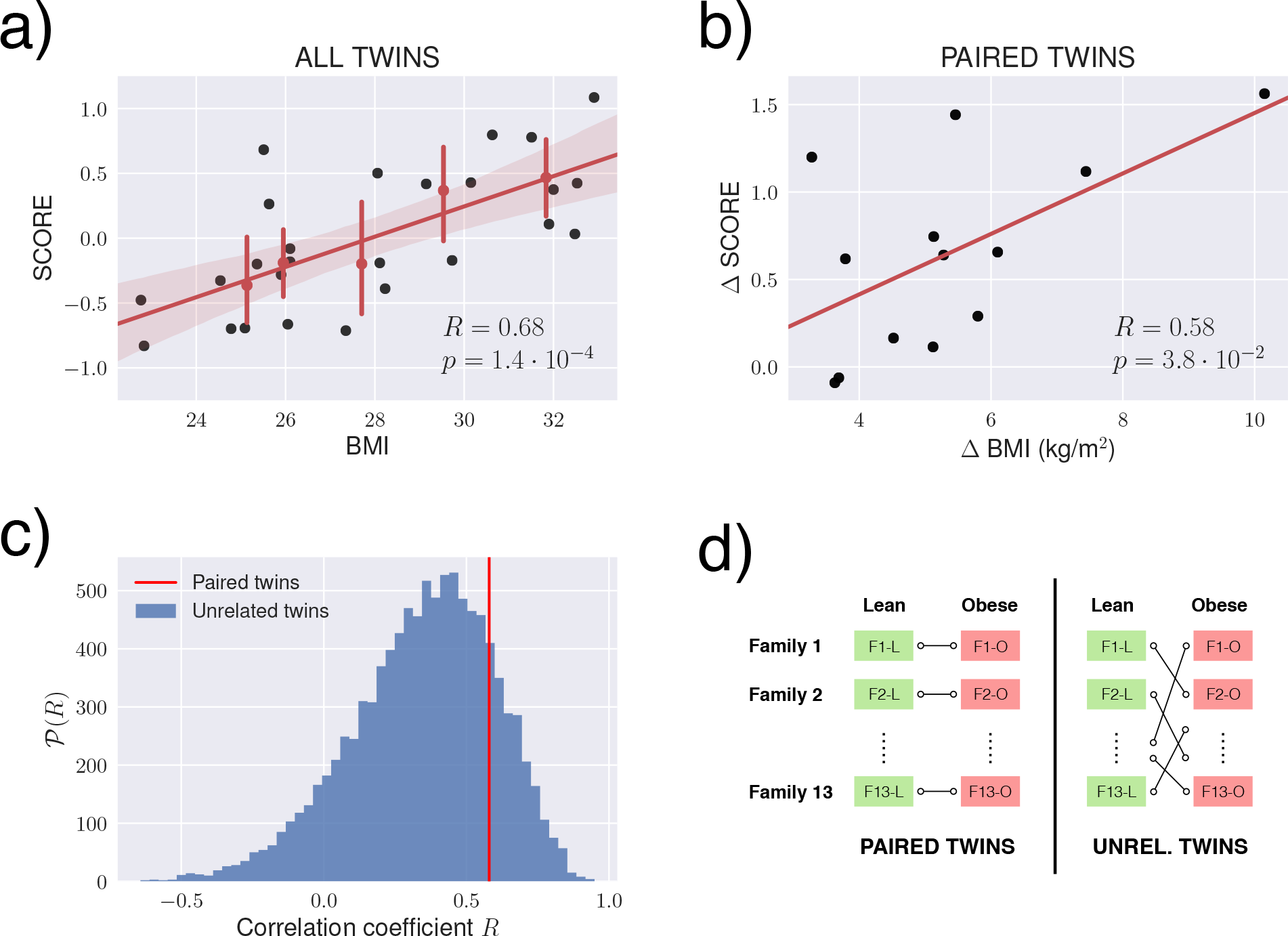
The obesity score in monozygotic twins. (a) Scatter plot of the obesity score versus BMI, for 26 adipose tissue samples corresponding to 13 pairs of monozygotic twins. (b) Scatter plot of change of obesity score versus change in BMI. Each point corresponds to a pair of twins, and change values are computed always as “obese co-twin versus lean co-twin”. (c) Comparison of the correlation coefficient R associated to paired twins with those obtained from random samples of unrelated twins, see (d).

## Conclusions

Rare genetic mutations in the leptin gene and elsewhere in the genome can cause extreme obesity, but the contribution of genetic over epigenetic and environmental factors in the current obesity pandemia is still debated. In this brief report, we show that obesity is correlated with a 38 genes transcriptomic signature in a BMI-dependent manner, even in subjects with the same genetic background. This suggests that pathway deregulation in obesity is linked to life-style rather than genes. Thus the only way to fight this disease is to work on the first aspect: If obesity is not due to the *bad luck* associated with inherited unlucky genes, each subject should in principle be able to reverse his/her condition by changing lifestyle.

## Methods

### Transcriptomic data

We use transcriptomes of 26 adipose tissue samples from 13 pairs of monozygotic twins with discordant BMI (BMI differences 3.3 – 10 kg/m^2^) from,^7^ which can be accessed under identifier E-MEXP-1425 at the ArrayExpress repository. In particular, we use the files labeled as “processed” and, after averaging out probes that map to the same gene, apply a simple normalization imposing that the mean gene expression is constant among samples.

### Obesity score

The obesity score *S_j_* for sample *j* is calculated as a linear combination of the log2 expression of 38 genes:

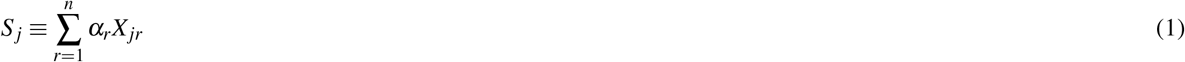

where *α*_*r*_ is the coefficient of the rank *r* gene in Table 1 and *X_jr_* is its log2 expression in sample *j*. The set of 38 genes and their coefficients shown in Table 1 were determined in^5^ using a dataset unrelated to the one analyzed in this manuscript. Obesity scores are displayed as mean-centered values in all figures.

**Table 1.** The 38 genes in the transcriptomic signature of obesity and their associated coefficients. Genes are ranked by the absolute value of their coefficient. Details of how these genes and coefficients were computed can be found in.^5^

| rank | Entrez ID | Gene Symbol | Coefficient | rank | Entrez ID | Gene Symbol | Coefficient |
| --- | --- | --- | --- | --- | --- | --- | --- |
| 1 | 1278 | COL1A2 | 0.131 | 20 | 7045 | TGFBI | 0.0569 |
| 2 | 80763 | SPX | −0.126 | 21 | 25878 | MXRA5 | 0.0558 |
| 3 | 761 | CA3 | −0.0889 | 22 | 2982 | GUCY1A3 | 0.0556 |
| 4 | 219348 | PLAC9 | 0.0742 | 23 | 2335 | FN1 | 0.0555 |
| 5 | 25975 | EGFL6 | 0.0731 | 24 | 7076 | TIMP1 | 0.0553 |
| 6 | 2014 | EMP3 | 0.0701 | 25 | 5396 | PRRX1 | 0.0548 |
| 7 | 6696 | SPP1 | 0.0690 | 26 | 4069 | LYZ | 0.0529 |
| 8 | 1397 | CRIP2 | 0.0679 | 27 | 8076 | MFAP5 | 0.0510 |
| 9 | 1490 | CTGF | 0.0674 | 28 | 3512 | JCHAIN | 0.0486 |
| 10 | 22822 | PHLDA1 | 0.0667 | 29 | 10402 | ST3GAL6 | −0.0466 |
| 11 | 1880 | GPR183 | 0.0659 | 30 | 3429 | IFI27 | 0.0458 |
| 12 | 171024 | SYNPO2 | 0.0655 | 31 | 83442 | SH3BGRL3 | 0.0457 |
| 13 | 1520 | CTSS | 0.0646 | 32 | 712 | C1QA | 0.0442 |
| 14 | 80114 | BICC1 | 0.0638 | 33 | 474344 | GIMAP6 | 0.0441 |
| 15 | 115207 | KCTD12 | 0.0622 | 34 | 9457 | FHL5 | 0.0438 |
| 16 | 151887 | CCDC80 | 0.0599 | 35 | 8470 | SORBS2 | 0.0437 |
| 17 | 22918 | CD93 | 0.0591 | 36 | 7037 | TFRC | 0.0431 |
| 18 | 389136 | VGLL3 | 0.0588 | 37 | 1291 | COL6A1 | 0.0430 |
| 19 | 8542 | APOL1 | 0.0581 | 38 | 57863 | CADM3 | 0.0429 |

## Acknowledgments

FFC and SZ are supported by the ERC advanced grant SIZEFFECTS. SZ acknowledges support from the Academy of Finland FiDiPro progam, project 13282993.

## Author contributions

FFC analyzed data. SZ and CAMLP designed the research and wrote the paper with the assistance of FFC.

